# Platelet concentrate-derived extracellular vesicles promote adult hippocampal neurogenesis

**DOI:** 10.1101/2025.04.29.651163

**Authors:** Ariunjargal NyamErdene, Nhi Thao Ngoc Le, Ouada Nebie, Emilie Faivre, Liling Delila, Ming-Li Chou, Joshua R Lowe, Kadiombo Bantubungi, Luc Buée, Tara L Walker, David Blum, Thierry Burnouf

## Abstract

Platelet-derived materials are emerging as promising, cell-free biotherapies for regenerative medicine. While platelet lysates have shown neuroprotective activity in preclinical models, the neurogenic potential of platelet concentrate-derived extracellular vesicles (pEVs) remains underexplored. Here, we evaluated the effects of human pEVs and a neuroprotective heat-treated human platelet lysate (HPPL) on adult hippocampal neurogenesis using both an ex vivo neurosphere assay and an in vivo intranasal administration model. pEVs selectively enhanced dentate gyrus (DG)-derived neurosphere growth, even in the absence of exogenous growth factors, and were internalized by neural precursors. In vivo, short-term pEV delivery increased EdU⁺ proliferating cells in the DG, while long-term administration (28 days) elevated the proportion of newborn mature neurons. By contrast, HPPL primarily promoted early neurogenesis by expanding immature DCX⁺ neurons. Quantitative proteomics of DG tissue after pEV treatment revealed 111 differentially expressed proteins, with enrichment in pathways related to oxidative phosphorylation, Notch4 signaling, myelination, and MHC class I-mediated antigen presentation. Downregulated proteins included cytoskeletal and translation-related regulators, suggesting a shift toward neuronal differentiation and circuit integration. Biophysical characterization confirmed the purity and vesicular nature of pEVs, with a defined protein cargo including immune modulators and ECM-interacting molecules such as CD44, lymphatic vessel endothelial hyaluronan receptor 1 (LYVE1), and complement proteins. These findings identify allogeneic pEVs as multifunctional agents that modulate neural precursor cell fate and brain tissue remodeling through coordinated metabolic and immunoregulatory mechanisms. This work supports the translational potential of pEV-based therapeutics for promoting hippocampal neurogenesis and cognitive repair in neurodegenerative and age-related brain disorders.

## 1. Introduction

Adult neurogenesis, the process by which neural stem/progenitor cells (NSPCs) generate new neurons and contribute to brain plasticity [1], continues throughout life in discrete brain regions, notably the subventricular zone (SVZ) and the subgranular zone (SGZ) of the hippocampal dentate gyrus (DG) [2, 3]. This process plays an essential role in cognition, memory formation, affective functions, and mood regulation [4, 5]. However, neurogenesis progressively declines with aging and is impaired in neurodegenerative and neuropsychiatric disorders such as Alzheimer’s disease and major depressive disorder [6, 7]. While physical activity [8, 9], environmental enrichment [10], and systemic trophic factors [11] can partially counteract this decline, there is a growing need for targeted, minimally invasive therapies to restore neurogenic capacity and neural circuit function [12].

Platelet-derived biomaterials are emerging as promising candidates for regenerative medicine owing to their inherent multifaceted bioactivity, abundance, and favorable safety profile [13]. Platelet concentrates (PCs), used routinely for transfusion, are a clinically available and scalable source of lysates and extracellular vesicles (EVs) that contain a rich array of growth factors, cytokines, antioxidants and anti-inflammatory factors [13]. Upon activation, platelets release extracellular vesicles (pEVs), nanoscale particles (50–200 nm) with a phospholipid bilayer that protects and transports cargo including proteins, nucleic acids, and metabolites [14]. These vesicles share some functional similarities and therapeutic potentials with mesenchymal stromal cell-derived EVs [15] and can be produced consistently under GMP-compliant conditions thanks to pooling strategies [13]. Several trophic factors abundant in platelet lysates and pEVs, such as brain-derived neurotrophic factor (BDNF), platelet-derived growth factor (PDGF), vascular endothelium growth factor (VEGF), epithelium growth factor (EGF), and platelet factor 4 (PF4; CXCL4) [16, 17] are individually known to exert neuroprotective effects in models of stroke [18], traumatic brain injury [19, 20] and neurodegeneration [21–24]. PF4, in particular, has recently been identified as a circulating exerkine that promotes hippocampal neurogenesis and improves cognitive function in aging models [9, 11]. These findings provide a strong biological rationale for investigating platelet-derived products as naturally enriched, multimodal agents for brain repair and neurorestoration [25–27]. pEVs can cross biological barriers and be taken up by recipient cells via endocytosis or membrane fusion, making them attractive for regenerative applications [13, 28]. Their capacity to modulate inflammation, angiogenesis, and oxidative stress has been demonstrated in cardiovascular [29], corneal injury models [30, 31] and wound healing [32]. Yet, the potential of native PC-derived EVs capable of stimulating adult hippocampal neurogenesis remains unexplored.

In this study, we systematically assessed the neurogenic effects of two platelet-derived products, heat-treated human platelet pellet lysate (HPPL) [19, 21] and PC-derived EVs on neural precursor cells in the DG. Using ex vivo neurosphere assays and in vivo intranasal delivery in adult mice, we evaluated their ability to stimulate NSPC proliferation, differentiation, and maturation. In parallel, we performed proteomic profiling of hippocampal tissue to elucidate the molecular pathways modulated by these treatments. By leveraging the natural regenerative cargo of platelet-derived vesicles and lysates, this work identifies novel, biocompatible biomaterials with potential for treating aging-related cognitive decline and neurodegenerative diseases through the enhancement of endogenous neurogenesis.

## 2. Material and Methods

### 2.1. Preparation and characterization of platelet materials

#### 2.1.1. Serum-converted platelet lysate (SCPL) and human platelet pellet lysate (HPPL)

Therapeutic-grade human PCs were collected from healthy volunteers at the Taipei Blood Center under Institutional Review Board approval (TMU-JIRB N201802052). These allogeneic PCs were suspended in 100% plasma, anticoagulated with citrate-phosphate dextrose solution A, and collected using the MCS+ system (Haemonetics, USA) and transported within 90 minutes under controlled temperatures to Taipei Medical University (TMU). PCs were stored on a slow-speed agitator at 22 ± 2 °C until processing within 24 hours. For this study, at least three PC units were initially pooled to generate both heat-treated platelet pellet lysate (HPPL) and serum-converted platelet lysate (SCPL), which was subsequently used for extracellular vesicle (EV) isolation and functional assays. All preparations were performed under aseptic conditions. To prepare SCPL, 0.3 mL of 0.9 M CaCl₂ and 5 g of sterile glass beads were added to every 10 mL of PC to induce coagulation and platelet activation. The mixture was rotated at room temperature until clot formation, followed by an additional 30-minutes rotation. Samples were centrifuged at 6,000 × g for 30 minutes, and the supernatant (SCPL) was collected, aliquoted, and stored at −80 °C until EV isolation.[33] To prepare HPPL, PCs were centrifuged at 3,000 × g for 30 minutes at 22 °C to pellet platelets. Plasma was removed, and platelet pellets were washed with PBS and concentrated to one-tenth of their original volume. The concentrated platelets underwent three freeze-thaw cycles (−80 ± 1 °C to 37 ± 1 °C), followed by centrifugation at 4,500 × g for 30 minutes at 20 ± 1 °C. The lysate was then heat-treated at 56 ± 1 °C for 30 minutes, cooled on ice for at least 5 minutes, and centrifuged at 10^4^ × g for 15 minutes at 4 ± 2 °C to remove precipitates.[21] For all experiments, HPPL was aliquoted (100 µL) in 1.5 mL Eppendorf tubes and stored at −80 °C until use.

#### 2.1.2. Isolation of platelet extracellular vesicles (pEVs)

Platelet concentrate-derived EVs (pEVs) were isolated and characterized as previously described [33]. Briefly, pEVs were purified from SCPL using size exclusion chromatography (SEC). A Merck Vantage L Laboratory column (VL 11 × 250 mm), packed with 10 mL of Sepharose CL-2B (Sigma-Aldrich, GE Healthcare Bio-Sciences, Sweden), was used for SEC with a flow rate of 1.2 mL/min. A 0.5 mL aliquot of SCPL was injected per run and eluted using PBS containing 0.32% sodium citrate (pH 7.4) as the running buffer [34]. pEV-enriched fractions were collected and pooled. Pooled fractions were concentrated using Amicon Ultra-4 centrifugal filter devices (50K cutoff, Merck Millipore) by centrifugation at 4,000 × g for 25 minutes. For experimental use, pEVs were aliquoted (100 µL) into 1.5 mL Eppendorf tubes and stored at −80 °C until use in ex vivo and in vivo assays.

#### 2.1.3. Biochemical and biophysical characterization of platelet materials

Total protein content in pEVs and HPPL was measured using a bicinchoninic acid (BCA) assay (Thermo Scientific, Cat# 23227). Particle concentration and size distribution were analyzed by nanoparticle tracking analysis (NTA) using a NanoSight NS300 system (Malvern), with samples diluted in 0.22 µm-filtered PBS to achieve 50–200 particles per frame. Each sample underwent five 60-second recordings, analyzed using NTA software (v3.4). In addition, dynamic light scattering (DLS) was used to assess size distribution using a Zetasizer Lab instrument (Malvern) and Zetasizer software (v7.13).

#### 2.1.4. Cryo-electron microscopy (cryo-EM) of SCPL and pEVs

Crude SCPL and SEC-purified EV samples were prepared for cryo-electron microscopy (cryo-EM) to assess vesicle morphology. A 4 µL aliquot of either SCPL-raw (obtained by serum conversion of platelet concentrates) or SEC-purified SCPL-EVs (resuspended in PBS-citrate used to equilibrate the SEC column) was applied to separate 200-mesh holey carbon grids. Grids were glow-discharged for 20 seconds using a plasma cleaner (Electron Microscopy Sciences) to improve surface hydrophilicity. Samples were blotted for 3 seconds under 100% humidity at 4 °C and immediately vitrified using a FEI Vitrobot Mark II. Grids were stored in liquid nitrogen until imaging. Micrographs were acquired at optimal magnification using a FEI Tecnai F20 transmission electron microscope, operated by staff at the Cryo-EM Core Facility, Academia Sinica, Taipei, Taiwan.

#### 2.1.5. Electrophoretic protein profiling of pEVs and HPPL

To visualize the protein composition of EVs and HPPL, sodium dodecyl sulfate–polyacrylamide gel electrophoresis (SDS-PAGE) was performed. A total of 20 µg of protein from each sample was mixed with XT sample buffer (Bio-Rad, Cat# 1610791) and heated at 95 °C for 5 minutes, either in the presence or absence of reducing agent (Invitrogen, Cat# NP0009). Samples were then loaded onto a precast NuPAGE 4–12% Bis-Tris protein gel (1.0 mm; Invitrogen, Cat# NP0321BOX). For reduced samples, antioxidant (Invitrogen, Cat# NP0005) was added to the running buffer according to the manufacturer’s protocol. Electrophoresis was initially run at 60 V for 30 minutes, followed by 125 V for the remainder of the run (Power supply: T-Pro Biotechnology, Model JB07-G003). After separation, gels were stained with InstantBlue Coomassie protein stain (Abcam, Cat# ab119211) for 15 minutes. Protein bands were visualized using a white-light transilluminator and documented with a digital camera.

#### 2.1.6. Detection of pEVs surface markers in EVs

A human-specific antibody membrane array (EXO-RAY210B-8, System Biosciences) was used to detect EV surface markers following the manufacturer’s instructions. EVs (50 µg total protein) were lysed in the provided buffer, and 1 µL of labeling reagent was added. Samples were incubated with continuous mixing for 30 minutes at room temperature. Excess labeling reagent was removed using the included separation column and wash steps. The labeled lysate was mixed with 5 mL of blocking buffer and applied to the antibody array membrane, followed by overnight incubation at 4 °C on a shaker. After washing, membranes were incubated with detection reagents for 30 minutes at room temperature. Streptavidin-HRP (5 µg/mL) and chemiluminescent substrates were then applied. Signal detection was performed using the UVP BioSpectrum 810 imaging system. Band intensities were quantified by subtracting background values from each target signal using the array software.

#### 2.1.7. Quantification of pEVs and platelet markers in EVs by nano flow cytometry (nFCM)

Surface marker expression of EV-specific and platelet-derived antigens in pEVs was analyzed using nano flow cytometry (nFCM). Samples were stained with fluorescently conjugated antibodies against CD9, and CD41followed by incubation at 37 °C. After staining, samples were washed with PBS and ultracentrifuged at 100,000 × g for 70 minutes at 4 °C. Pellets were resuspended in PBS for subsequent analysis. To ensure optimal detection, samples were diluted to achieve particle acquisition rates between 2,000 and 12,000 events per minute. Particle size distribution, concentration, and surface marker expression were assessed using a NanoFCM flow cytometer. Data were processed with NanoFCM Profession software (version 2.330), with vesicle size and concentration calculated from flow rate and side scatter intensity, using standard calibration curves.

### 2.2. Ex vivo neurosphere assay

#### 2.2.1. Animals

Eight-week-old adult female C57BL/6J mice (n = 15) were obtained from the Taiwan National Laboratory Animal Center (Nangang, Taipei, Taiwan) and used for neural precursor cell isolation. All procedures were conducted in accordance with institutional ethical guidelines and approved by the Institutional Animal Care and Use Committee of TMU (Protocol No. SHLAC2023-0050)

#### 2.2.2 Neural precursor cell culture

Neural precursor cells were isolated from the dentate gyrus (DG) of the hippocampus of 8-week-old female C57BL/6J mice, as previously described [35]. DG tissues were dissected and processed separately. DG tissue was minced and enzymatically digested at 37 °C for 14 minutes: DG samples with papain, with intermittent gentle agitation. Following digestion, DG cells were dissociated mechanically by gentle pipetting. Cell suspensions were incubated for an additional 10 minutes, centrifuged at 300 × g for 5 minutes, and resuspended in hank’s balanced salt solution (HBSS). The suspensions were passed through a 40 μm cell strainer (Corning, Cat No. 431750) to remove debris. After final centrifugation, cell pellets were resuspended in 200 μL of complete growth medium composed of Neurobasal medium [Gibco #21103-049], with 2% B-27 [Gibco #17504-044], 1% penicillin/streptomycin (10,000 U/mL), [Gibco #15140122], 1% GlutaMAX [Fisher Scientific #35050061], and 20 ng/mL each of EGF [Sigma #AF-100-15] and FGF-2 [Sigma #100-18B]). In some experiments, EGF and FGF-2 were not included and replaced by pEVs or HPPL. For proliferation assays, cells were cultured with 0.5% pEVs, 0.5% HPPL, or no supplementation (control). Neurosphere formation was assessed after 10 days.

#### 2.2.3. Proliferation assay

The proliferation assay was performed as previously described [35]. Briefly, neural precursor cells from DG were cultured in proliferation medium supplemented with either 0.5% pEVs or 0.5% HPPL. Control wells received no supplementation. After 10 days of culture, proliferation was assessed by counting the number of neurospheres in 80 wells of a 96-well plate per condition and measuring their diameter of 50 randomly selected neurospheres using a light microscope (Leica). Additionally, to evaluate the proliferative effects of platelet-derived materials in the absence of exogenous mitogens, a separate set of three independent experiments was performed using medium lacking EGF and FGF-2. In these assays, 0.5% pEVs or HPPL was added as the sole supplement, and unsupplemented medium served as a negative control. All measurements were performed in a blinded manner to avoid bias, with experiments repeated independently five times (n = 5).

#### 2.2.4. Alexa fluor 488 Labeling of pEVs and HPPL and their internalization in neurospheres

pEVs and HPPL were covalently labeled with Alexa Fluor 488 using the Protein Labeling Kit (A10238, Thermo Fisher Scientific) per manufacturer’s instructions. Samples were resuspended in 0.1 M sodium bicarbonate buffer (pH 8.3) at 1 mg/mL, combined with reconstituted Alexa Fluor 488 dye in a 1:10 protein ratio, and incubated overnight at 4 °C in darkness with gentle agitation. Labeled products were purified using the provided columns pre-equilibrated with elution buffer, washed with PBS to remove unbound dye, and stored at 4 °C for up to two weeks. Labeling efficiency was confirmed by measuring absorbance/emission at 555/565 nm. For internalization, neurospheres from DG (10-day culture) were grown in low-adhesion plates with supplemented neurobasal medium, then incubated with labeled pEVs and HPPL (10-50 µg/mL) for 24 hours at 37 °C with 5% CO₂. After PBS washing, neurospheres were transferred to coated coverslips, fixed with 20% cold methanol, stained with DAPI (1:5000), and imaged using Stellaris microscope with Z-stack to visualize 3D internalization within the neurospheres.

### 2.3. In vivo neurogenesis

The primary animal study was conducted at the ULille Animal Research Center (Inserm), and a replication animal study was performed at the Taipei Medical University (TMU) Shuangho animal facility. Mice at both facilities were housed under standard conditions (12-hour light/dark cycle; temperature 18.5–24 °C; humidity 40–60%). Animals were allowed to acclimate for one week prior to the start of experiments. All procedures were conducted in accordance with institutional ethical guidelines and approved by the Institutional Animal Care and Use Committee of ULille and TMU (APAFIS 43474-2023050714441306 and SHLAC2023-0050).

#### 2.3.1. Intranasal (i.n) administration of pEVs and HPPL for short-term neuronal proliferation and long-term maturation analysis

pEVs, HPPL, and PBS vehicle (control) were prepared in 100 μL aliquots and thawed immediately prior to use. A total volume of 30 μL was administered intranasally (i.n.) to 8-week-old female C57BL/6J mice (n = 18 per group) while mice were held in an upright position. The administration followed a precise protocol where small drops (2-3 μL each) were delivered alternately to each nostril with 1-2 minutes intervals between drops until the total dose of 30 μL was reached. For short-term proliferation analysis, mice received daily intranasal administration for three consecutive days (total of 3 doses). For long-term neurogenesis assessment, a separate cohort received intranasal administration every 3^rd^ day until D27 and sacrificed at D28 days (total of 11 doses).

#### 2.3.2. 5-Ethynyl-2^′^-Deoxyuridine (EDU) injection

To label proliferating neural precursor cells for both short-term and long-term neurogenesis assesments, mice received six intraperitoneal (i.p.) injections of 5-ethynyl-2′-deoxyuridine (EdU; 50 mg/kg; Sigma, 900584), a thymidine analog that incorporates into DNA during the S-phase of the cell cycle [36]. Injections were administered twice daily (T0 and T6 hours) for three consecutive days starting 2 hours after the first i.n. administration of pEVs, HPPL, or PBS vehicle. EdU was freshly prepared in sterile, filtered PBS and dissolved immediately prior to injection. Body weights were monitored at baseline (week 0), week 2, and week 4 to assess general health during the study period.

#### 2.3.3. Histological procedures

Mice were deeply anesthetized using a combination of Zoletil 50 (50 mg/mL; Sigma) and Xylazine (23.32 mg/mL; Sigma), diluted in double-distilled water. Transcardial perfusion was performed with cold 0.9% NaCl, followed by 4% paraformaldehyde (PFA) in 0.1 M phosphate buffer (pH 7.4). Brains were dissected and post-fixed in 4% PFA for 24 hours at 4 °C, then cryoprotected in 30% sucrose (JT Baker) for 48 hours. Following cryoprotection, brains were immersed in pre-cooled isopentane (–35 to –40 °C) for 1 minute and stored at –80 °C until sectioning. Coronal sections (35 μm) were collected for proliferation analysis, and sagittal sections (35 μm) from one hemisphere were used for neuronal maturation studies. Sections were obtained using a Leica CM3050S cryostat (TMU core facility), transported on dry ice, and stored in PBS containing 0.2% (w/v) sodium azide at 4 °C until use.

#### 2.3.4. Immunofluorescence protocol

Free-floating brain sections spaced 350 µm apart, spanning the dorsal dentate gyrus to the lateral ventricles (Bregma –1.34 to –2.70 mm), were collected in ten series per animal. Sections were washed in cold PBS and permeabilized in PBS containing 0.2–0.5% Triton X-100 for 30 minutes with gentle agitation. For EdU staining, slices were blocked in 3% BSA in PBS for 20 minutes, followed by a second permeabilization with 0.5% Triton X-100 for 20 minutes. After an additional 20-minutes blocking step, sections were incubated with the EdU Click-iT cocktail (Thermo Fisher) for 30 minutes at room temperature. Sections were then rinsed three times in PBS with 3% BSA and counterstained with DAPI (1:500; Thermo Fisher, 62248) for 5 minutes. For co-staining with neuronal markers, slices were incubated in a mouse-on-mouse blocking reagent (Vector MKB2213, 1:100 in 0.2% Triton X-100 PBS) for 1 hour. Primary antibody staining was performed with mouse anti-NeuN-biotinylated (Millipore MAB377B; 1:500 in 1% normal goat serum) and anti-DCX (1/250, Millipore Ab2253, LOT2881137) for 48 hours at 4 °C. Following PBS washes, sections were incubated with Streptavidin-Alexa Fluor 568 (Invitrogen S11226; 1:500) for 1–2 hours at room temperature. After staining, sections were mounted on adhesive slides and air-dried in the dark. Autofluorescence was quenched by sequential treatment with 70% ethanol (5 minutes), 0.1% Sudan Black B in 70% ethanol (5 minutes), and a final 70% ethanol rinse (1 minute). Slides were washed in 0.9% NaCl and coverslipped using DAKO fluorescent mounting medium (DAKO S3023).

#### 2.3.5. Image acquisition

Brain tissues from C57BL/6J mice (n = 18 per group) were processed for immunofluorescence and stained with EdU and/or NeuN. Brain sections were scanned using the TissueFAXS slide scanning system (ZEISS, model 1025727155) at 20× magnification (120 slides total), and representative images were captured using Stellaris confocal microscopy with 63× objectives. Each animal contributed 3–5 comparable coronal brain sections for analysis. Quantification of positively stained cells was performed using HistoQuest software (TissueGnostics, version 5). Fluorescence intensities were analyzed across individual channels, and events were recorded as counts per region. Analyses were focused on the hippocampal DG, corresponding to Bregma coordinates –1.34 to –2.70 mm, based on the Allen Brain Atlas.

#### 2.3.6. Sample processing for proteomics

For pEV and HPPL samples, protein precipitation was performed using cold acetone. Acetone was pre-cooled to −20°C for at least 24 hours prior to sample preparation. The pEV and HPPL samples were mixed with cold acetone at a 1:4 ratio. After vortexing, the samples were incubated overnight at −20°C. The next day, samples were centrifuged at 15,000 × g for 10 minutes at 4°C. The supernatant was discarded, and the pellets were washed twice with cold acetone/water (4:1), followed by centrifugation at 15,000 × g for 10 minutes after each wash. Pellets were air-dried on ice and then resuspended in 6 M urea (prepared by dissolving 1.8 g of urea in 5 mL of water). Protein concentrations were measured using the BCA assay, and the final concentration was adjusted to at least 1 μg/μL. Approximately 100 μg of protein was submitted to the National Taiwan University (NTU) Proteomics Core Facility for LC-MS/MS analysis. In-solution digestion was performed at NTU. Briefly, proteins were reduced with 5 mM DTT at 29°C for 45 minutes, then alkylated with 10 mM iodoacetamide in the dark at 29°C for another 45 minutes. Trypsin was added at a 1:50 (enzyme, w/w) ratio for overnight digestion (16 h) at 29°C. Digestion was stopped by adding 10% TFA to a final concentration of 0.5%. Samples were desalted using C18 columns prior to LC-MS/MS analysis

#### 2.3.7. Mouse tissues of dentate gyrus (DG) following pEVs and PBS treatment

DG tissues were collected at 28 days post-intranasal administration (dpi) from mice treated with PBS (n = 3) and pEVs (n = 3). Proteins were homogenized using ice-cold 10 mM Tris base (pH 7.4) containing 10% sucrose (0.32 M). Samples were sonicated on ice at 10% amplitude (10 sec on/5 seconds off, total 1 minute), maintaining temperature below 30°C. Lysates were homogenized for 1 hour at 4°C using a rotor. After centrifugation at 15,000 × g for 10 minutes, supernatants were collected, and protein concentration was immediately determined by BCA analysis.

For LC-MS/MS, 200 µg of total protein was precipitated with cold acetone (1:4, v/v) and incubated overnight at –20°C. Samples were centrifuged (15,000 × g, 10 min, 4°C), washed twice with cold acetone, air-dried, and resuspended in 6 M urea. Following re-quantification, tissues from 3 mice per group were pooled to obtain 100 µg of protein per sample. In-solution digestion was performed at NTU, where proteins were reduced with 5 mM DTT at 29°C for 45 minutes, then alkylated with 10 mM iodoacetamide in the dark at 29°C for another 45 minutes. Trypsin was added at a 1:50 (enzyme, w/w) ratio for overnight digestion (16 h) at 29°C. Digestion was stopped by adding 10% TFA to a final concentration of 0.5%. Samples were desalted using C18 columns prior to LC-MS/MS analysis. Fresh samples were submitted to the National Taiwan University (NTU) Proteomics Core Facility for LC-MS/MS using an Orbitrap Fusion Lumos Tribrid quadrupole-ion trap-Orbitrap mass spectrometer (Thermo Fisher Scientific).

#### 2.3.8. Bioinformatic analysis of proteomics data

Raw data were analyzed using the Mascot search engine integrated in Proteome Discoverer software (version 2.2, Thermo Scientific). Searches were conducted against the most recent UniProt (Swiss-Prot) mouse protein database (release: 2023), with parameters including: trypsin as the digestion enzyme, maximum of two missed cleavages, precursor mass tolerance of 10 ppm, and fragment mass tolerance of 0.02 Da. Carbamidomethylation of cysteine was set as a fixed modification, while oxidation of methionine and N-terminal acetylation were set as variable modifications. Peptide-spectrum matches were filtered at both protein and peptide levels using a target-decoy strategy. For protein identification, high-confidence peptide-spectrum matches (PSMs) were filtered using a false discovery rate (FDR) of 1%, with additional filters including “IsMasterProtein” and Mascot score ≥ 30. Only proteins quantified in both technical replicates were included in the analysis. Differentially expressed proteins (DEPs) were identified using a two-tailed unpaired t-test assuming unequal variances (heteroscedastic). Proteins with a fold-change ≥ 1.2 or ≤ 0.83 and a p-value < 0.05 were considered significantly differentially expressed. Gene Ontology (GO) enrichment analysis was conducted using DAVID Bioinformatics Resources 6.8, while canonical pathways and upstream regulators were further analyzed with Ingenuity Pathway Analysis (IPA, Qiagen).

### 2.4. Statistical analysis

All data are presented as mean ± standard deviation (SD) or standard error of the mean (SEM), as indicated in figure legends. Statistical analyses were performed using GraphPad Prism (version 10.2.3; GraphPad Software, La Jolla, CA, USA). Differences among the pEVs, HPPL, and control groups were assessed using ordinary one-way ANOVA followed by Dunnett’s multiple comparisons test to compare treatment groups against the control. A *p* value < 0.05 was considered statistically significant. The number of independent biological replicates for each experiment is indicated in the corresponding figure legends.

## 3. Results

### 3.1. Characterization of platelet-derived extracellular vesicles (pEV): comparative analysis with heat-treated pellet platelet lysates (HPPL) for neurogenesis application

We isolated extracellular vesicles (EVs) from serum-converted platelet lysates using size exclusion chromatography with Sepharose CL-2B resin, following established protocols [37]. Both pEVs and HPPL were generated from pooled platelet concentrates (n=3) obtained from Taipei Blood Center (Fig. 1A). To confirm the specific characteristics of pEVs, we first characterized the samples based on their morphology. Cryo-electron microscopy confirmed the presence of membrane-bound vesicles in pEVs preparations, displaying typical cup-shaped morphology with sizes around 100 nm (Fig. 1B). For comparison, cryo-EM images of the raw materials (serum-converted platelet lysates) shown in (Fig. 1B-left) revealed an unclear lipid layer with a dirty background surrounded by other lipids, in contrast to the isolated EVs with their distinct membrane structures.

**Fig.1.**
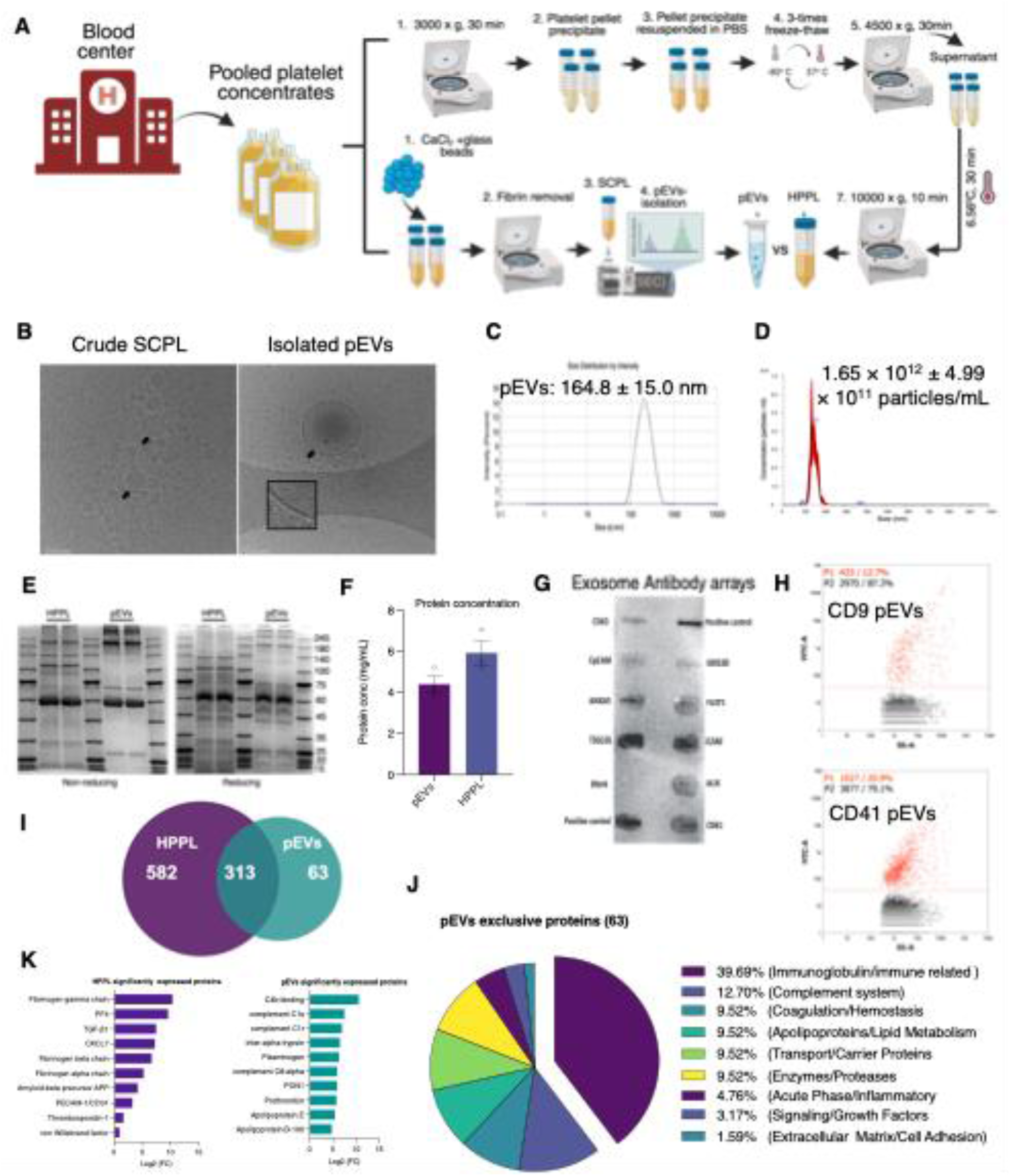
Comprehensive characterization of platelet-derived extracellular vesicles (pEV) and HPPL isolated from clinical-grade platelet concentrates. **(a)** Schematic workflow for isolation and preparation of pEVs and HPPL from clinical-grade platelet concentrate. (**b**) Representative cryo-electron microscopy (cryo-EM) images showing EVs from SCPL (left) and SEC-isolated EVs-SCPL (right). Arrows indicate characteristic double-membrane structure. Scale bar = 100 nm. (**c**) Size distribution profiles of pEVs determined by dynamic light scattering (DLS) across three independent batches (mean ± SD, n = 3). (**d**) Quantification of total particle yields from pEVs as measured by nanoparticle tracking analysis (NTA). (**e**) SDS-PAGE analysis of pEVs and HPPL under reducing and non-reducing conditions (20 μg protein/lane). Molecular weight markers are shown in the rightmost lane. (**f**) Protein concentration are shown by BCA (n=3) (**g**) Exo-Check antibody array demonstrating expression of canonical EV markers (CD63, Tsg101, and CD81) in pEVs. (**h**) Nano flow cytometric analysis of CD9 and CD41 expression in pEVs at optimized particle counts (2,000-12,000 particles/min), with size distribution and concentration determined through software calibration. (**i**) Venn diagram illustrating the proteomic profiles of pEVs and HPPL, showing the number of shared and unique proteins identified in each preparation. (**j**) Functional categorization of proteins exclusively detected in pEVs, expressed as percentage distribution across categories. (**k**) Top 10 significantly enriched proteins in pEVs and HPPL, ranked by expression ratio and quantified as log2 fold change (Log2FC) of pEVs versus HPPL.

The morphological observations were further validated by size distribution analysis. Dynamic light scattering revealed that pEVs exhibited a uniform size distribution with a mean diameter of 164.8 nm (Fig. 1C), consistent with the sizes observed in our cryo-EM analysis. Complementing these findings, nanoparticle tracking analysis demonstrated that pEVs contained significantly higher particle concentrations (1.65 × 10¹² particles/mL) (Fig. 1D). Having established the basic physical and morphological characteristics of our isolated pEVs, we selected heat-treated pellet platelet lysates (HPPL) as our comparative control for subsequent analyses. To comprehensively compare these two platelet-derived preparations, we proceeded to analyze their protein profiles, concentrations, and molecular signatures as detailed below.

We first examined the general protein profiles in both samples by SDS-PAGE. Analysis under non-reducing and reducing conditions revealed distinct protein profiles between pEVs and HPPL (Fig. 1E). Both conditions showed prominent bands around 60-65 kDa, likely corresponding to protein disulfide isomerase or albumin in both samples. Under reducing conditions, additional bands appeared: approximately 245 kDa in pEVs and 100-140 kDa in HPPL, possibly representing integrin subunits. These distinct banding patterns suggest the presence of disulfide-dependent protein complexes that are differentially distributed between these preparations. Total protein quantification by BCA revealed comparable concentrations between pEVs (4.4 mg/mL) and HPPL (5.9 mg/mL) preparations (n=3, P > 0.05; Fig. 1F), validating our subsequent comparative protein profile analysis.

Having established the physical and basic protein characteristics of our preparations, we proceeded to analyze specific markers to confirm the vesicular nature of our isolates. Surface marker analysis using Exo-check demonstrated the presence of canonical EV markers in pEVs, including tetraspanins (CD63, CD81), endosomal sorting proteins (TSG101), and membrane-associated protein FLOT1 (Fig. 1G low cytometry further confirmed the expression of CD9 (12.7%), a typical EV surface membrane marker, and platelet-specific marker CD41 (20.9%) (Fig. 1H), verifying that the pEVs retained markers related to their platelet origin.

To gain deeper insights into the molecular composition of our preparations, we also conducted comprehensive proteomic analysis. LC-MS/MS analysis of the pEVs revealed a proteome distinct from that of HPPL. Among the identified proteins, 313 were shared between the two preparations, while 63 were unique to pEVs and 582 to HPPL (Fig. 1I). Notably, approximately 39% of the pEV-specific proteins were immunoglobulins and about 12% were complement-related proteins (Fig. 1J). The shared protein fraction between pEVs and HPPL also showed significant differences in protein expression levels (Fig. 1K), with HPPL significantly expressing proteins such as platelet-specific PF4, TGF-beta1, and CXCL7. Particularly significant for potential therapeutic applications, approximately 1% of the pEV-specific proteome consisted of extracellular matrix (ECM) and cell adhesion molecules (Fig. 1K), including CD44, LYVE1, and HABP2 (Table S1).

### 3.2. pEV selectively enhance neural precursor cell proliferation in the dentate gyrus

To explore how pEVs and HPPL influence neural precursor cells, we focused on neurospheres isolated from dentate gyrus (DG) of 8-week-old adult female mice. Using pooled cells from three mice per region helped us minimize individual variations (Fig. 2A). Our initial investigation examined whether supplementing the medium with pEVs and HPPL (0.5% v/v) could influence neurosphere development. Interestingly, we found that pEVs specifically increased the size of DG neurospheres without affecting their number (Fig. 2B). To further validate these distinct effects, we examined neurosphere formation in the absence of exogenous growth factors. Notably, the specific responses persisted under these stringent conditions, with pEVs continuing to selectively enhance DG-derived neurosphere size (Fig. 2C). To assess treatment uptake by neurosphere cells, we exposed them to Alexa Fluor 488-labeled pEVs and HPPL for 24 hours. Confocal microscopy revealed progressive accumulation of fluorescent signal within neurospheres throughout this period, demonstrating that neural progenitor cells efficiently internalize both pEVs and HPPL (Fig. S1).

**Fig.2.**
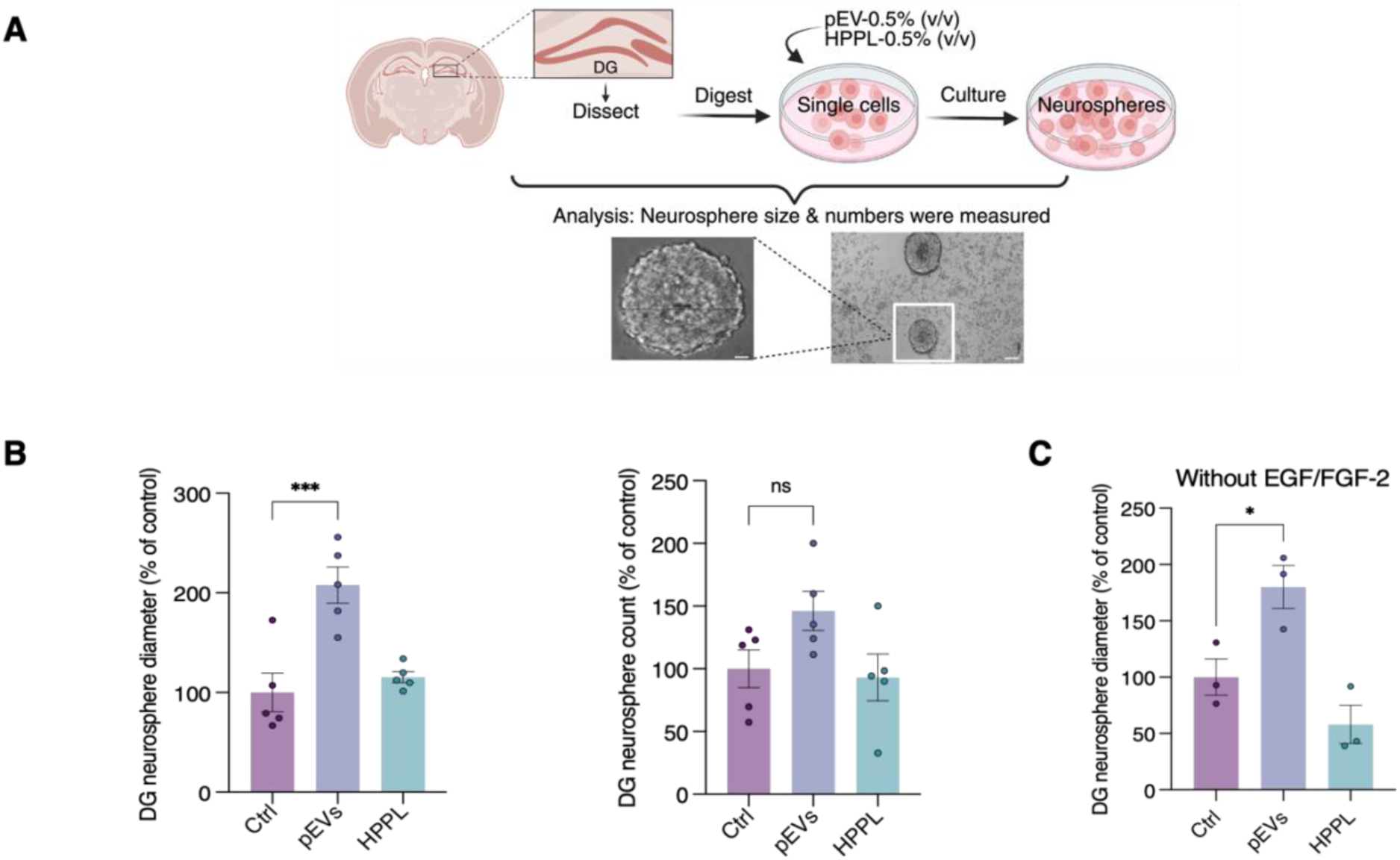
pEVs selectively regulate DG-derived neural precursor cell proliferation *ex vivo*. **A.** Schematic representation of neural stem cell isolation from the dentate gyrus (DG) followed by neurosphere generation. **B**. Quantification of neurosphere number and size distribution from DG-derived cultures following treatment with platelet concentrate-derived extracellular vesicles (pEVs) or heat-treated platelet pellet lysate (HPPL). The size of 50 neurospheres was analyzed per condition. For neurosphere number, all visible neurospheres were counted in 80 wells of a 96-well plates per condition. Data are mean ± SEM from five independent experiments. ***p < 0.001 vs PBS using one-way ANOVA followed by Dunnett’s multiple comparisons post-hoc test. Control condition is normalized to 100%, and that pEVs and HPPL values represent the relative change in neurosphere size and number compared to this baseline. **C.** Proliferation of DG-derived neurospheres treated with pEVs or HPPL in the absence of EGF and FGF-2. Data represent mean ± SEM from three independent experiments. *p* < 0.05 by one-way ANOVA with multiple comparisons.

To corroborate our ex vivo findings, we tested pEVs-specific functions in the DG of hippocampus in vivo. We administered pEVs, HPPL, or PBS (control) intranasally for three consecutive days, along with daily EdU injections administered six hours apart from the intranasal treatments to track newly dividing proliferating cells (Fig. 3A). Mice were sacrificed on day 4, and DG hippocampal tissue sections were analyzed for EdU staining. Quantification revealed significantly higher numbers of proliferating cells along the granular layer in pEVs-treated animals compared to controls (Fig. 3B), substantiating our ex vivo observations. These findings confirm that pEVs specifically enhance proliferation in the DG of the hippocampus.

**Fig.3.**
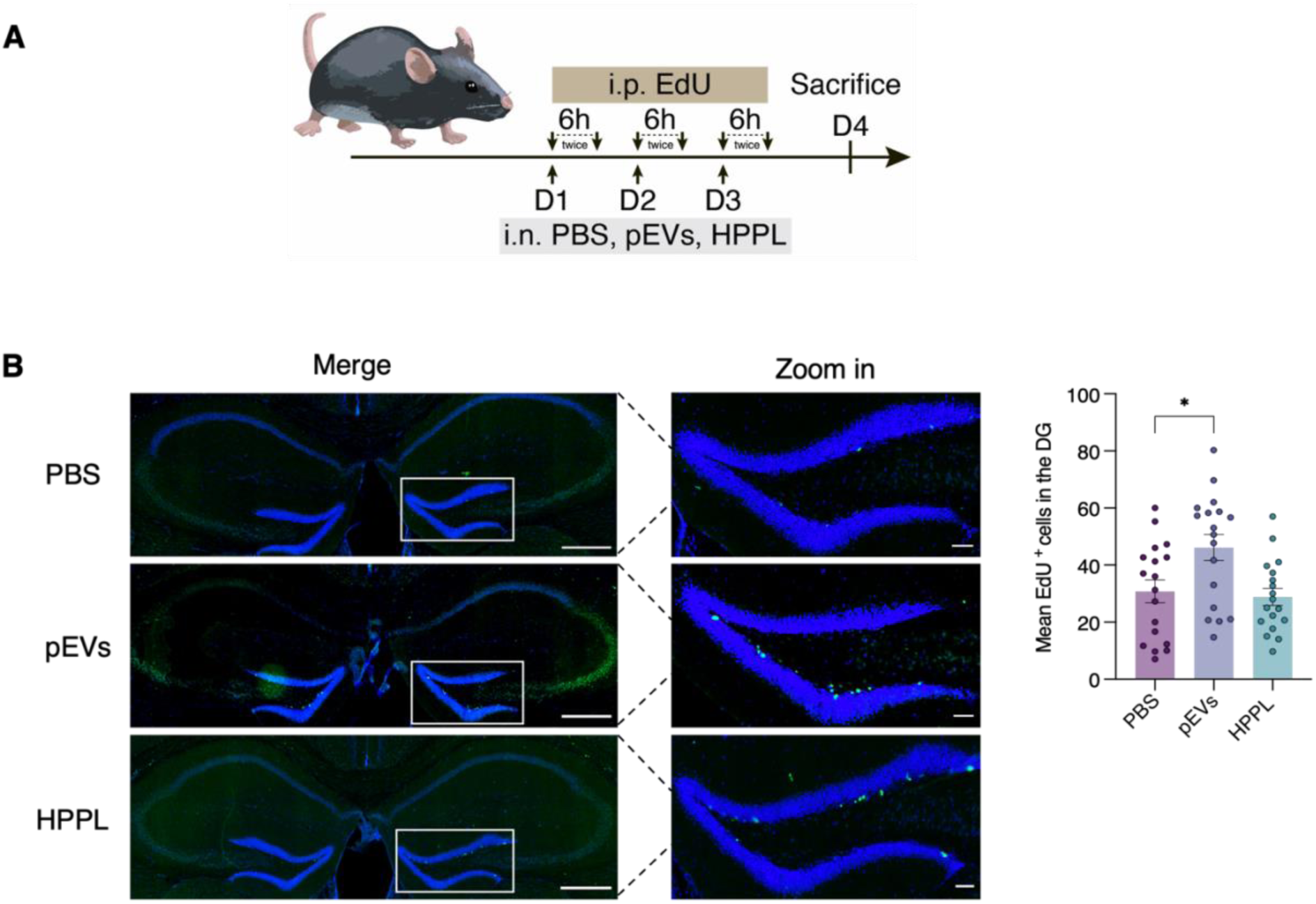
pEVs selectively regulate DG neural precursor cell proliferation *in vivo* at day 3. **A.** Schematic representation of the short-term (3-day) intranasal delivery protocol for pEVs or HPPL, combined with EdU (50 mg/kg) i.p administration twice daily at 6-hour intervals. **B.** Representative confocal images of EdU⁺ cells in the DG following 3-day treatment with pEVs or HPPL. Green channel represents EdU-labeled cells, and blue channel shows DAPI nuclear staining. Images acquired by Stellaris confocal microscopy with (20× objectives). Scale bar: 100 μm (inset: 50 μm). Quantification of total EdU^+^ cells across 5 DG sections/animal along the granular layer in DG. Data are mean ± SEM (N = 18 animals per group). *p < 0.05 vs PBS using one-way ANOVA followed by Dunnett’s multiple comparisons post-hoc test.

### 3.3. pEVs and HPPL Exert Divergent Effects on Neural Precursor Cell Differentiation

To further analyze how pEVs influence the cell fate of ex vivo neuronal precursor cells, we evaluated differentiation patterns of second-passage neurospheres from the dentate gyrus treated with pEVs or HPPL (0.5% v/v) for 10 days (Fig. 4A). At the end of the incubation period, we stained the DG-derived neurospheres with markers for immature neurons (DCX^+^) and mature neurons (NeuN^+^). Interestingly, HPPL treatments significantly promoted differentiation into immature DCX^+^ neuroblasts (Fig. 4B-C). In contrast, pEVs treatment significantly induced differentiation of neural precursor cells into mature NeuN^+^ neurons rather than immature DCX^+^ neuroblasts (Fig. 4B-D).

**Fig.4.**
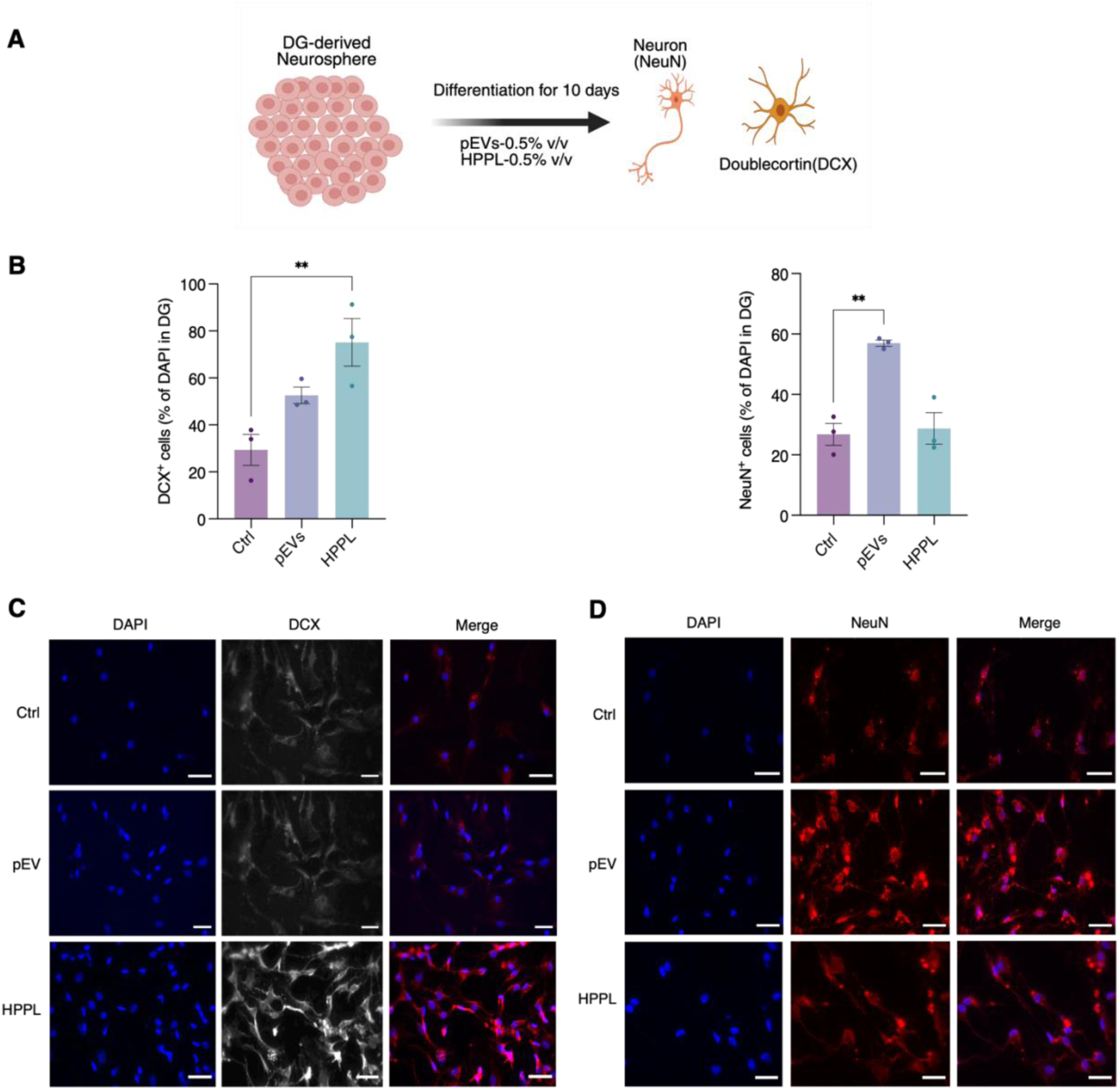
Differentiation of neural precursor cells ex vivo under pEV and HPPL stimulation. **A.** Schematic representation of the experimental design for neurosphere differentiation studies under pEVs and HPPL conditions. **B.** Quantification of DCX, and NeuN-positive cells as a percentage of total DAPI-positive nuclei per field. Data represent mean ± SEM from (n=3) ordinary one-way ANOVA followed by Dunnett’s (**p < 0.01) with multiple comparisons post-hoc test. **C.** Representative immunocytochemistry images showing DCX expression (red) in differentiated neurospheres under pEVs or HPPL treatment compared to untreated controls. Nuclei counterstained with DAPI (blue) Scale bars = 50μm **D.** Representative immunocytochemistry images of DG-derived differentiated neurospheres immunostained for NeuN (red) following pEV or HPPL treatment versus untreated controls. Nuclei counterstained with DAPI (blue) Scale bars = 50μm.

### 3.4. Long-term assessment of hippocampal neurogenesis in the dentate gyrus under pEV

To investigate the long-term neurogenic effects of platelet-derived extracellular vesicles (pEVs) and heat-treated platelet pellet lysates (HPPL), a separate cohort of mice received intranasal administration of either pEVs or HPPL three times per week over a 28-day period. To trace dividing cells, 5-ethynyl-2′-deoxyuridine (EdU) was administered during the first three days of treatment (Fig. 5A). This strategy allowed the identification of cells that entered the cell cycle early during the treatment window.

**Fig.5.**
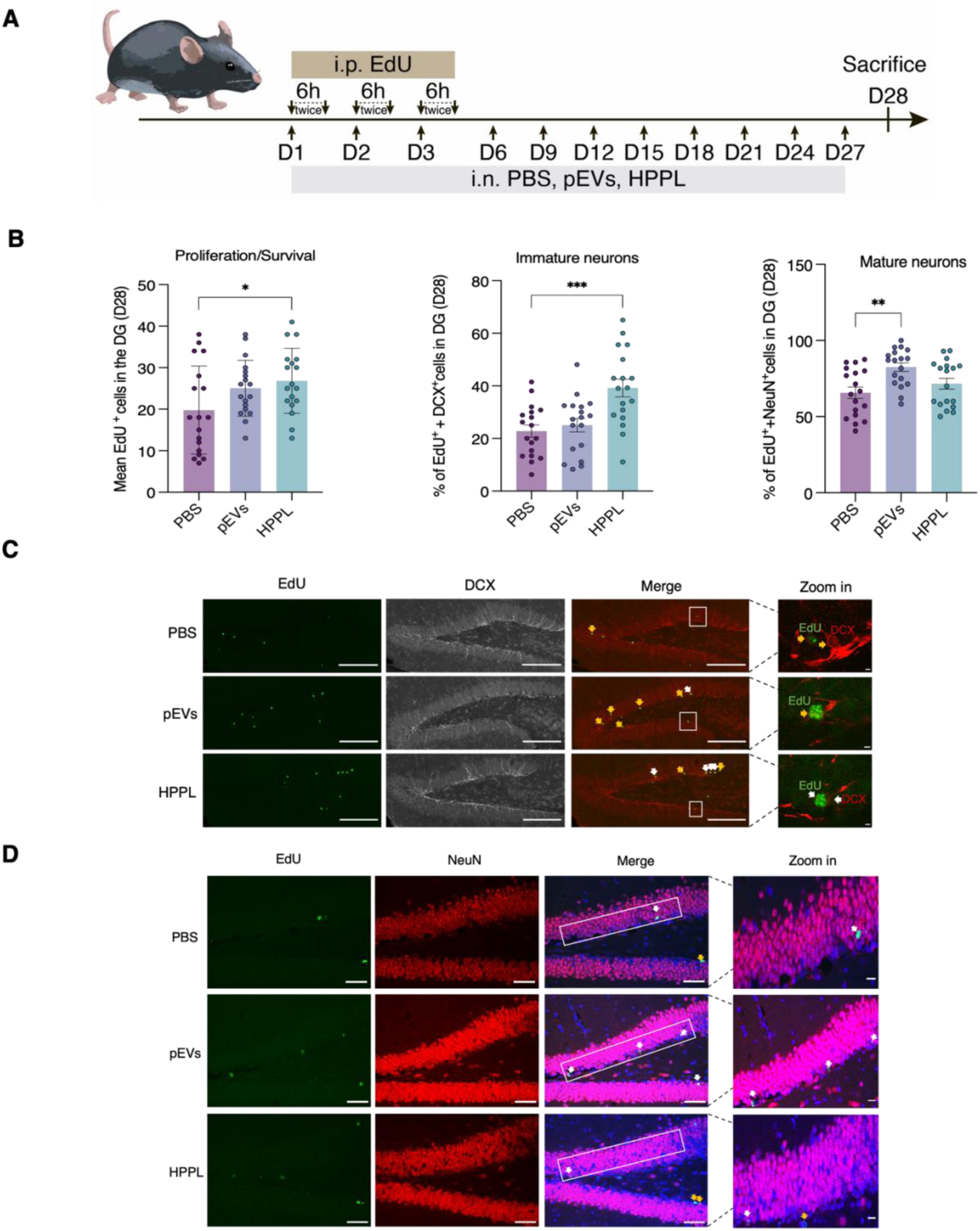
pEVs promote neurogenesis in the dentate gyrus (DG) of hippocampus. **A.** Experimental design schematic showing long-term (28-day) intranasal administration protocol for pEVs and HPPL combined with EdU (50 mg/kg) labeling regimen (twice daily at 6-hour intervals). **B.** Quantification shows total EdU^+^ cells across 5 DG sections/animal along the granular layer in DG, percentage of co-expressing DCX^+^/EdU^+^ and NeuN^+^/EdU^+^ cells in the granular layer of DG region. Data represent mean ± SEM (N = 18 animals per group). Statistical significance determined by ordinary one-way ANOVA followed by Dunnett’s (*p < 0.05, **p < 0.01 \*\*\**p < 0.001) with multiple comparisons post-hoc test.* **C-D.** Representative images of immature/mature neurons co-expressing DCX^+^/EdU^+^ and NeuN^+^/EdU^+^ in the granular layer of DG following pEVs & HPPL treatment at day 28. DCX is illustrated by gray channel, EdU by green channel, and NeuN, by red channel. Scale bars: 50 μm; inset scale bar: 20 μm. White arrows indicate DCX^+^/EdU^+^, and NeuN^+^/EdU^+^ co-expressing cells, orange arrows indicate DCX^-^/EdU^+^, and NeuN^-^/EdU^+^ cells. Representative images were captured using Stellaris confocal microscopy with (63× objectives). Data indicate that pEVs significantly increase the proportion of NeuN⁺ cells among EdU⁺ cells in the DG, suggesting enhanced neuronal differentiation, while HPPL primarily promotes cell proliferation and increases the number of immature neurons in the DG.

At the end of the 28-day treatment, immunohistochemistry revealed a significant increase in the total number of EdU⁺ cells in the dentate gyrus (DG) of HPPL-treated mice (Fig. 5B-C), suggesting enhanced cellular proliferation and/or survival. Moreover, a notable increase in the proportion of EdU⁺ cells co-expressing doublecortin (DCX), a marker of immature neurons, was observed in the HPPL group. This indicates that HPPL not only boosts proliferation but may also promote early neurogenic differentiation.

While pEV-treated mice showed a trend toward an increase in total EdU⁺ cells, this change was not statistically significant (Fig. 5B-D). However, pEV treatment significantly elevated the proportion of EdU⁺ cells that co-expressed NeuN, a marker of mature neurons, within the DG granule cell layer. This suggests that pEVs may facilitate the maturation and survival of newborn neurons over time.

### 3.5. Proteomic Insights into pEV-mediated modulation of dentate gyrus neurogenesis

To unveil the mechanisms underlying the effects of pEVs on dentate gyrus of hippocampal neurogenesis, we performed a proteomic analysis of DG tissue following 28 days of intranasal pEV or PBS treatment (N = 2 per group; Fig. 6A). 2,656 proteins were identified in the pEV group versus 2,629 in the PBS group, with 2,580 proteins (95.4%) shared. 76 proteins were unique to pEV-treated tissue, and 49 to PBS (Fig. 6B). GO-term analysis of the 76 pEV-specific proteins in DG tissue showed enrichment in protein binding and cell adhesion molecule binding functions (Table S2; Fig. S2). Differential proteomic analysis identified 111 significantly regulated proteins, with 61 upregulated and 50 downregulated (Fig. 6C). The upregulated proteins were primarily associated with biological processess of CNS myelination, oxygen binding, and postsynaptic signaling, and were enriched in oxidative phosphorylation and neuroplasticity-related KEEG pathways (Fig. 6D, red). The most profoundly altered proteins following pEV treatment are detailed in (Table S3). Among the significantly upregulated proteins, several key regulators of neurogenesis emerged with angiotensin-converting enzyme (ACE; Log_2_FC: 1.6537), which suggests altered regulation of the renin-angiotensin system. Structural neuronal proteins were also elevated, such as internexin neuronal intermediate filament protein Alpha (INA; Log_2_FC: 1.0899) and developmentally regulated GTP-binding Protein 1 (DRG1; Log2FC: 1.2182). Additionally, proteins associated with neuronal maturation, including Ras-related protein Rab-7a (RAB7; Log_2_FC: 0.917) and brain abundant membrane-attached signal protein 1 (BASP1; Log_2_FC: 0.7989) were upregulated (Fig. 6D). In contrast, downregulated proteins were predominantly enriched in biological processes related to cytoplasmic translation, actin filament organization, and cell migration (Fig. 6D, blue). They included key cytoskeletal regulators such as Rho-associated coiled-coil containing protein kinase 1 (ROCK1; Log2FC: -0.7107), known for its role in actin dynamics and cell proliferation [38], as well as stathmin 1 (STMN1), a regulator of microtubule stability, and Ras-related C3 botulinum toxin substrate 1 (RAC1), a GTPase involved in lamellipodia formation. The reduced expression of these proteins suggests diminished migratory activity as neurons settle into their final positions. Ingenuity Pathway Analysis (IPA) confirmed activation of respiratory electron transport, myelination, and ubiquitination pathways (Fig. 6E). IPA also uniquely identified activation of Notch4 signaling, critical for neural stem cell maintenance [39] and MHC class I antigen presentation, suggesting coordinated neurogenic and immune-related responses following pEV treatment. Our molecular analyses revealed distinct biological pathways affected by the treatments. Expanded protein-protein interaction (PPI) analysis further corroborated these findings, with upregulated proteins distinctly clustering in myelination and proteasome pathways, while downregulated proteins were predominantly associated with translation processes. The comprehensive network visualization and analysis of these interactions is presented in (Fig. 7A-B), providing a systems-level perspective that integrates our findings.

**Fig.6.**
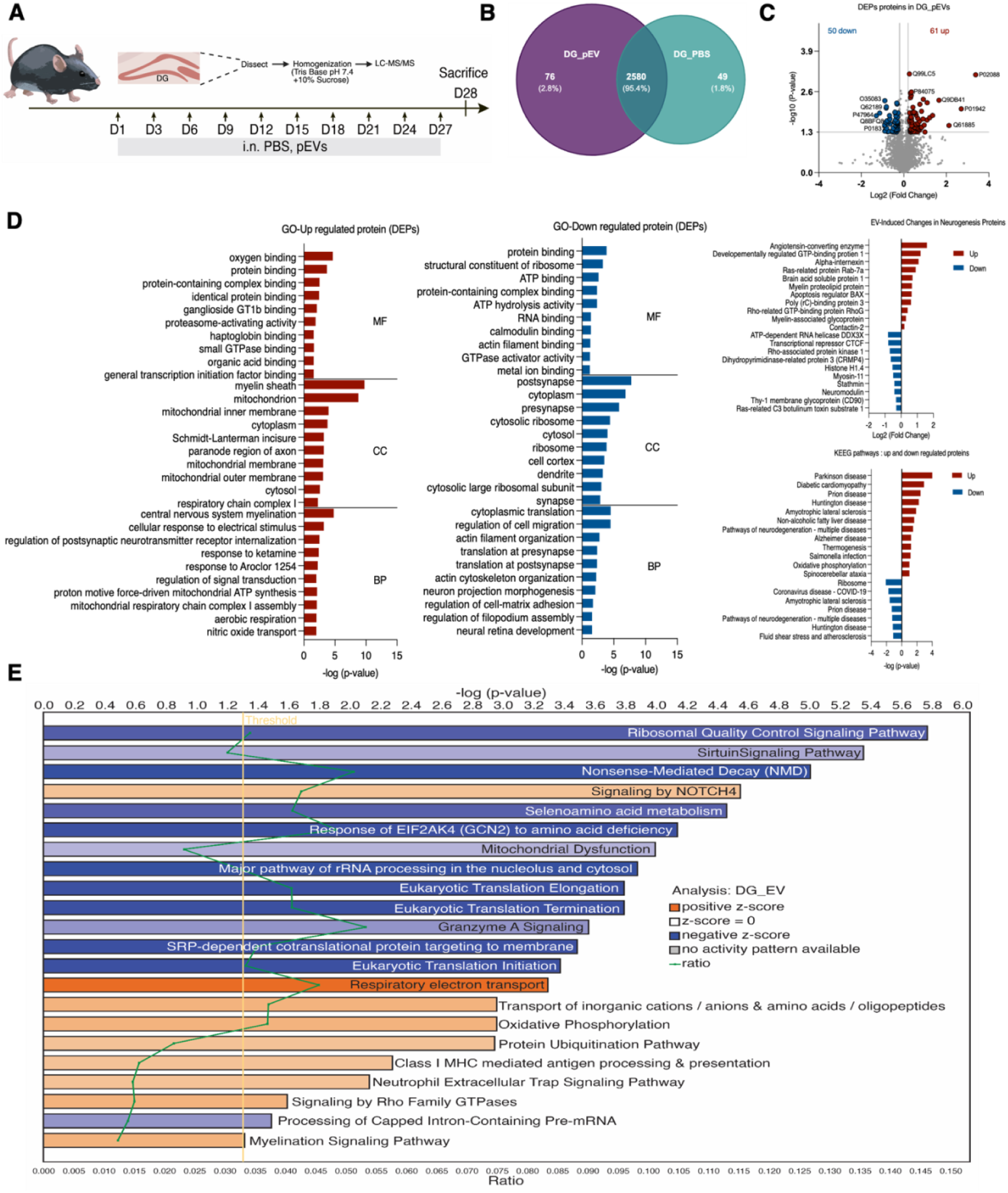
Proteomic profiling reveals pEVs-induced molecular changes in the dentate gyrus (DG) of hippocampus. **A**. Experimental workflow showing DG tissue collection following 28-day of intranasal administration of pEVs or PBS as control, followed by proteomic profiling via LC-MS/MS. **B.** Venn diagram illustrating the overlap of identified proteins in DG tissue from pEVs and PBS-treated groups, highlighting both shared and treatment-specific proteins. **C.** The differentially expressed proteins (DEPs) were identified by comparing proteins commonly expressed between pEV-treated samples and PBS controls. Volcano plot analysis of these DEPs in DG tissue following pEV treatment revealed significant changes using thresholds of p ≤ 0.05 and fold change ≥ 1.2. Red and blue points on the plot indicate significantly upregulated and downregulated proteins, respectively. **D.** Gene Ontology (GO) enrichment of upregulated and downregulated DEPs categorized by biological process, molecular function, and cellular component. Neurogenesis Pathway Proteins Altered by EV Treatment: Upregulated and Downregulated Proteins with Log2FC Values, KEGG pathway analysis is shown for both subsets. **E.** Canonical pathway analysis of differentially expressed proteins (DEPs) using Ingenuity Pathway Analysis (IPA), highlighting significantly altered signaling networks (p < 0.05) after DG_EV treatment compared to PBS. Pathways are color-coded by activation z-score: orange bars indicate predicted activation (positive z-score), blue bars indicate predicted inhibition (negative z-score), and white bars indicate pathways with z-score = 0. The green line represents the ratio of DEPs present in each pathway relative to the total proteins in that pathway.

**Fig.7.**
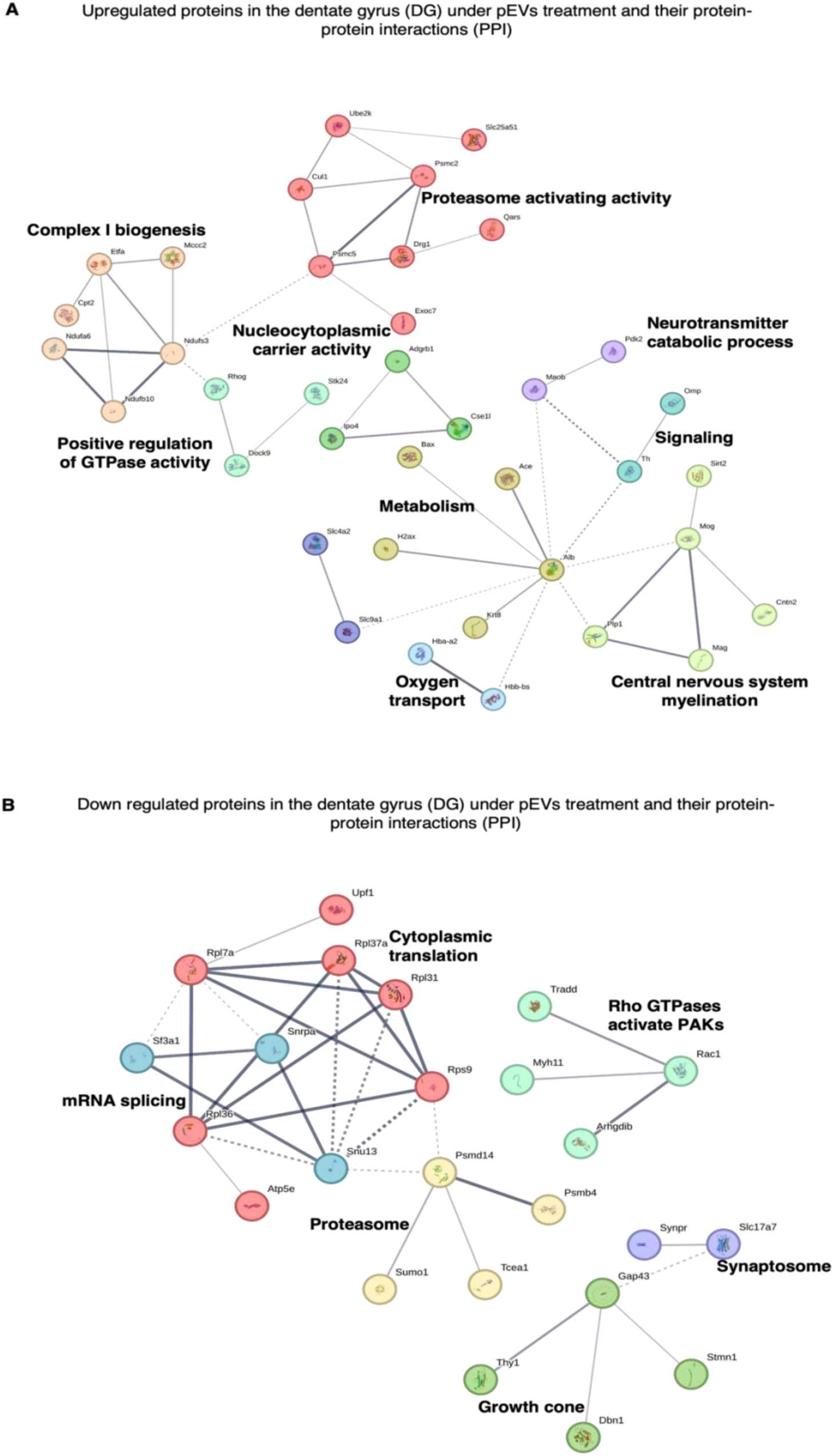
STRING protein-protein interaction network analysis of upregulated and downregulated proteins in DG following pEVs treatment. (a) Network was generated using MCL clustering algorithm with inflation parameter of 3 to identify natural clusters based on stochastic flow. Network statistics: 61 nodes (proteins), 45 edges (interactions), average node degree of 1.48, and average local clustering coefficient of 0.423. The network shows significant enrichment in protein-protein interactions (PPI enrichment p-value: 0.000114) compared to expected interactions (24). Visualization parameters: maximum FDR ≤ 0.05, minimum signal ≥ 0.01, and minimum interaction strength ≥ 0.01. (b) Network was generated using MCL clustering algorithm with inflation parameter of 3 to identify natural clusters based on stochastic flow. Network statistics: 50 nodes (proteins), 34 edges (interactions), average node degree of 1.36, and average local clustering coefficient of 0.344. The network shows significant enrichment in protein-protein interactions (PPI enrichment p-value: 0.02) compared to expected interactions (23). Visualization parameters: maximum FDR ≤ 0.05, minimum signal ≥ 0.01, and minimum interaction strength ≥ 0.01.

## 4. Discussion

This study provides new insights into the use of pEVs as bioactive, cell-free materials able to modulate adult hippocampal neurogenesis. Using ex vivo neurosphere cultures and in vivo intranasal delivery in adult mice, we demonstrate that pEVs selectively enhance proliferation and maturation of DG neural precursor cells. Our proteomic analysis reveals that pEVs modulate hippocampal protein expression in pathways related to mitochondrial function, myelination, and immune signaling. These findings support the translational potential of pEVs as a next-generation neurorestorative therapy for age-related and neurodegenerative brain disorders.

Previous studies have demonstrated the neuroprotective potential of HPLs administered via intranasal route in rodent models of Parkinson’s disease [21, 23] and Alzheimer’s disease [24], via intracerebroventricular injection in models of amyotrophic lateral sclerosis (ALS) [22], and directly applied to the brain surface in models of traumatic brain injury (TBI) [20]. These beneficial effects have been linked to the rich content of trophic factors (e.g. BDNF, VEGF, PDGF, PF4) [21, 22] and functional biomolecules such as antioxidants [40, 41], which protect against various forms of cell death and oxidative stress. HPLs consistently outperform standard recombinant growth factors, with their protective effects involving Akt pathway activation and context-dependent involvement of MEK signaling [42]. pEVs represent a more defined fraction enriched in membrane-enclosed cargo and free from large plasma proteins, likely enabling better mechanistic clarity, and, possibly, reproducibility [14, 43]. The current study builds on this rationale by directly comparing pEVs with a benchmark neuroprotective lysate, HPPL [21].

The DG of the adult hippocampus is a key site of lifelong neurogenesis in adults, providing holistic roles in learning and memory [44] but is highly sensitive to aging and disease [45]. Our ex vivo and in vivo findings show that, as compared to HPPL, pEVs enhance proliferation of DG-derived neural progenitors ex vivo, even in the absence of EGF and FGF-2, suggesting intrinsic trophic effects. Neural progenitor cell fate also differed markedly between treatments with both platelet biomaterials. Indeed, pEVs directed differentiation toward mature NeuN^+^ neurons, while HPPL only promoted development of immature neuronal cells in both neurosphere ex vivo analysis and in vivo. Overall, our data demonstrate that pEVs exert a powerful impact on DG neurogenesis, significantly influencing both proliferation and differentiation processes. This comprehensive effect underscores the potential of pEVs as modulators of adult hippocampal neurogenesis across multiple stages of neural development

Interestingly, our DG proteomic finding revealed the complex molecular landscape underlying pEV-mediated neurogenic effects in the dentate gyrus, suggesting a sophisticated interplay between neurodevelopmental regulators and immunomodulator factors. The upregulation of key neurogenic regulators in DG tissue including ACE (modulating angiotensin signaling for progenitor proliferation)[46], INA and BASP1 (orchestrating cytoskeletal organization and transcriptional reprogramming) [47], DRG1 (coordinating GTPase activity for neuronal migration) [48], and RAB7 (optimizing endosomal trafficking for synaptogenesis) [49] reveals that pEV treatment induces a sophisticated protein expression profile that simultaneously influences multiple critical neurogenic processes from neural stem cell activation through terminal differentiation, ultimately favoring the development of mature NeuN^+^ neurons over immature neuroblasts. Simultaneously, the downregulation of cytoskeletal restrictors including ROCK1 (inhibitor of neurite outgrowth and branching)[50], RAC1 (controller of actin cytoskeleton dynamics) [51], and STMN1 (microtubule destabilizing protein) [52]. While these proteins are typically essential for various aspects of neuronal development, their downregulated expression in this context occurring alongside upregulation of key neurogenic factors would indicate a coordinated remodeling of the cytoskeletal regulatory network. This appears to create a precisely balanced neurogenic environment in response to our treatment, rather than simple inhibition of these pathways.

Further supporting these findings, IPA analysis revealed activation of Notch4 and MHC class I-related pathways, which is consistent with emerging evidence that Notch signaling plays a dual role in adult neurogenesis by maintaining the neural stem cell pool while directing lineage commitment toward neuronal fates [53, 54]. The concurrent activation of MHC class I-related pathways suggests pEVs create an immune-privileged microenvironment that would protect developing neurons during their vulnerable integration period [55–57]. This molecular signature represents a sophisticated regulatory network that balances stem cell maintenance with neuronal differentiation while simultaneously modulating neuroinflammatory processes known to influence the neurogenic niche. Collectively, these complementary mechanisms establish the molecular foundation underlying pEV-mediated neurogenic enhancement, particularly explaining the preferential generation of mature NeuN^+^ neurons observed consistently across our ex vivo and in vivo analyses. The EV proteome also included ECM-binding proteins (e.g., CD44, LYVE1, HABP2), which are known to interact with hyaluronan, a key ECM component known to regulate neural stem cell maintenance and proliferation, suggesting potential mechanisms for the biological effects of pEVs in neural tissue applications [58].

Immunoregulatory components (e.g., complement proteins, Ig fragments) were also detected in the EV proteome pointing to a potential role in remodeling the neurogenic niche. These immune-related factors may contribute to the creation of a permissive microenvironment by modulating local inflammatory processes [59], clearing cellular debris, and regulating microglial activation states. The complement proteins present in our EVs, particularly C4b, may engage with complement receptors on neural cells [60], potentially influencing cellular recognition, synaptic refinement, and microglial activity in ways that ultimately promote neurogenesis [61]. This finding adds to the growing evidence that immune-related molecules play non-canonical roles in neural development and plasticity, extending beyond their traditional immune functions [62]. Additionally, the identified immunomodulatory factors could regulate the cross-talk between neural and immune cells within the neurogenic niche [63], potentially suppressing detrimental pro-inflammatory signals while enhancing trophic support [64].

The pEVs used in this study were isolated from serum-converted platelet lysates using similar GMP-compatible SEC method as previously described in a cardiac ischemia/reperfusion [29]. In the cardiac ischemia study, SCPL-EVs improved cardiac function, reduced apoptosis and scarring, and promoted angiogenesis, with no adverse coagulation or systemic toxicity effects [29].

These data confirm the functional activity and safety of SCPL-EVs across tissues. Proteomic features such as the enrichment of HSP70, RAP1B, and LGALS3BP, and the depletion of coagulation-related proteins, were consistent across this and our study. Additional support for the translational versatility of pEVs comes from their successful use in ocular models [13]. A separate study from our group demonstrated that pEVs, isolated from PC supernatants by ultracentrifugation, efficiently delivered the anti-angiogenic agent kaempferol to the cornea, suppressed pathological neovascularization, and extended drug retention on the ocular surface [31] while also exerting its own beneficial biological effects in this application. This work highlighted the intrinsic anti-inflammatory and targeting properties of pEVs, and their capacity to modulate angiogenic and inflammatory gene expression in vascular endothelial cells. The ability of native pEVs to penetrate tissue barriers and enhance therapeutic retention further reinforces their relevance as delivery vectors for CNS therapies, where barrier crossing and local retention are critical [13, 27]. Furthermore, previous investigations from our group using fluorescently labeled HPPL and pEVs demonstrated that proteins from these materials can diffuse into the brain and reach the hippocampus following intranasal administration in mice [19, 40]. These findings therefore support the feasibility of delivering bioactive platelet-derived molecules, including those packaged in pEVs, to central brain regions through the intranasal route. Additional evidence for the therapeutic value of SCPL-EVs also comes from a recent study demonstrating their successful integration into gelatin-based hydrogel foams for wound healing applications [32]. In this model, the same type of pEVs as used in the current manuscript was incorporated into engineered hydrogels and applied to chronic wounds in diabetic rats. The formulation not only accelerated wound closure but also modulated macrophage polarization, promoted angiogenesis, and maintained a low-inflammatory environment [32]. These effects were consistent with the anti-inflammatory and pro-regenerative activity observed in the brain and reinforce the multifunctional capacity of pEVs to modulate both immune and tissue repair responses. The biocompatibility and safety of pEVs in this context further support their translation toward clinical applications in tissue regeneration, including CNS repair. Together, these findings support the use of pEVs as a well-characterized, reproducible, and scalable biomaterial for brain repair. Importantly, while this study emphasizes the distinct advantages of pEVs, it also confirms the functional activity of HPPL, which significantly enhanced the pool of immature neural progenitors both ex vivo and in vivo. This complementary effect suggests that HPPL may support early stages of neurogenesis, while pEVs promote later maturation. These observations raise the possibility that a combined therapeutic approach, leveraging both HPPL and pEVs, could provide additive or synergistic benefits across multiple stages of neural regeneration. Future studies should explore this hypothesis experimentally to optimize platelet-derived therapies for CNS repair. The consistent efficacy of PC-derived EVs in both cardiac and neural injury models, and promising performance in ocular delivery, highlights their broad regenerative potential and encourages further exploration for clinical applications in neurological diseases.

In conclusion, the ability of pEVs to modulate neuronal maturation and engage immune and metabolic pathways opens new avenues for non-cell-based interventions in neurodegenerative disease. Their compatibility with minimally invasive delivery (e.g., intranasal) and scalable production from clinical-grade PC [13] makes them attractive for translational development. Future work should assess long-term behavioral outcomes, explore their efficacy in aging and disease models, and dissect the contribution of specific vesicular components to neurogenesis and synaptic plasticity. Overall, this study advances the concept that extracellular vesicles derived from human platelet concentrates constitute a novel and versatile neuroregenerative platform with potential clinical utility.

## Supporting information

Complete Supplemental file

## CRediT authorship contribution statement

AN performed the experiments, analyzed the data and drafted the manuscript. LNTN conducted the proteomics analysis. ON performed part of the intranasal administrations at Lille University. EF assisted with the analysis of the brain sections. LD assisted with intranasal administration and mouse sacrifice procedures at TMU. MC assisted in proteomics data interpretation and design of figures. JL performed parallel ex vivo neurosphere assays using platelet lysates and discussed data. KB suggested and provided recommendations for the ex vivo neurosphere assays. LB contributed to methodological recommendations and funding. TLW provided expert recommendations all along the study, conducted data analysis, and drafted the manuscript. DB and TB are PhD co-advisors of AN, conceived the study design, provided funding support, and drafted the manuscript. All authors revised and approved the final version.

## Data availability

Data will be made available on request.

## Acknowledgments

We thank the Taipei Blood Center (Guangdu, Taiwan) for supplying the platelet concentrates and the Consortia of Key Technologies and Instrumentation Center at National Taiwan University for providing mass spectrometry technical services for brain tissue analysis.

## Appendix A. Supplementary data

