## Supplementary material for "Platelet concentrate-derived extracellular vesicles promote adult hippocampal neurogenesis": Complete Supplemental file

[1]International Ph.D. Program in Biomedical Engineering, College of Biomedical Engineering, Taipei Medical University, Taipei ,Taiwan, [2]University of Lille, Inserm, CHU Lille, UMR-S1172 Lille Neuroscience & Cognition (LilNCog), Lille, France, [3]Alzheimer and Tauopathies, LabEx DISTALZ, France, [4]Graduate Institute of Biomedical Materials and Tissue Engineering, Taipei Medical University , Taipei, Taiwan, [5]Clem Jones Centre for Ageing Dementia Research, Queensland Brain Institute, The University of Queensland, Brisbane, Australia, [6] Inserm UMR-1011, University of Lille, Lille, France, [7]International Ph.D. Program in Cell Therapy and Regeneration Medicine, [8]Ph.D. Program in Graduate Institute of Mind Brain and Consciousness, College of Humanities and Social Sciences, Taipei Medical University , Taipei , Taiwan.

**Contents:**

**Supplemental Tables (Table S1-S3)**
**Supplemental figures (Figure S1-S2)**

### Supplemental Tables (Tables S1-S3)

**Supplemental Table S1. Unique protein signatures associated with pEVs treatment identified by LC-MS/MS.** Proteins exclusively expressed in the pEVs group, as identified by LC-MS/MS, in comparison to the HPPL group.

| EVs-63 unique protein list | |
|---|---|
| **Accession** | **Protein Name** |
| **P01742** | Immunoglobulin heavy variable 1-69 |
| **A0A075B6S5** | Immunoglobulin kappa variable 1-33 |
| **P02747** | Complement C1q subcomponent subunit C |
| **P13671** | Complement component C6 |
| **Q96E17** | Ras-responsive element-binding protein 1 |
| **Q9Y5Y7** | Lymphatic vessel endothelial hyaluronic acid receptor 1 (LYVE1) |
| **P01718** | Immunoglobulin lambda variable 3-27 |
| **P55056** | Apolipoprotein C-IV |
| **Q15485** | Ficolin-2 |
| **P26927** | Hepatocyte growth factor-like protein |
| **P23083** | Immunoglobulin heavy variable 1-2 |
| **P15169** | Carboxypeptidase N catalytic chain |
| **P18428** | Lipopolysaccharide-binding protein |
| **P20851** | C4b-binding protein beta chain |
| **P04430** | Immunoglobulin kappa variable 1-16 |
| **P11226** | Mannose-binding protein C |
| **P0DTE1** | Immunoglobulin heavy variable 3-38 |
| **P01877** | Immunoglobulin heavy constant alpha 2 |
| **Q6UX71** | Plasma kallikrein-sensitive glycoprotein |
| **Q9NZP8** | Complement C1r subcomponent-like protein |
| **Q13790** | Apolipoprotein F |
| **A0A075B6K4** | Immunoglobulin kappa variable 1-5 |
| **P00740** | Coagulation factor IX |
| **P0DJI8** | Serum amyloid A-1 protein |
| **A0A0B4J1U3** | Immunoglobulin heavy variable 6-1 |
| **Q6UXB8** | Peptidase inhibitor 16 |
| **P54108** | Cysteine-rich secretory protein 3 |
| **P15814** | Immunoglobulin lambda-like polypeptide 1 |
| **A0A0C4DH34** | Immunoglobulin heavy variable 3-23 |
| **O14791** | Apolipoprotein L1 |

| | |
|---|---|
| **P22352** | Glutathione peroxidase 3 |
| **P11597** | Cholesteryl ester transfer protein |
| **P01704** | Immunoglobulin lambda variable 2-14 |
| **P01833** | Polymeric immunoglobulin receptor |
| **Q9UK55** | Protein Z-dependent protease inhibitor |
| **P05160** | Coagulation factor XIII B chain |
| **P0CF74** | Immunoglobulin lambda constant 6 |
| **P27918** | Properdin |
| **Q04756** | Hepatocyte growth factor activator |
| **P16070** | CD44 antigen |
| **P04180** | Phosphatidylcholine-sterol acyltransferase |
| **P0DOX3** | Immunoglobulin kappa constant |
| **P03952** | Plasma kallikrein |
| **P23142** | Fibulin-1 |
| **Q9BXR6** | Complement factor H-related protein 5 |
| **P01601** | Immunoglobulin kappa variable 1D-33 |
| **Q14520** | Hyaluronan-binding protein 2 |
| **P20742** | Pregnancy zone protein |
| **A0A0C4DH43** | Immunoglobulin kappa variable 2D-28 |
| **Q15166** | Paraoxonase 3 |
| **Q9Y6R7** | IgGFc-binding protein |
| **P35542** | Serum amyloid A-4 protein |
| **O00391** | Sulfhydryl oxidase 1 |
| **P29622** | Kallistatin |
| **P01714** | Immunoglobulin lambda variable 3-19 |
| **P02786** | Transferrin receptor protein 1 |
| **P02745** | Complement C1q subcomponent subunit A |
| **P01703** | Immunoglobulin lambda variable 1-40 |
| **A0A087WSX0** | Immunoglobulin lambda variable 5-45 |
| **P01767** | Immunoglobulin heavy variable 3-53 |
| **A0A0B4J1V6** | Immunoglobulin heavy variable 3-21 |
| **P02748** | Complement component C9 |
| **Q08380** | Galectin-3-binding protein |

**Supplemental Table S2. Functional Classification of Unique Proteins Identified in the DG Following pEVs or PBS Treatment.** Proteins identified in the DG following pEVs and PBS control treatments were analyzed using Venny 2.1. A total of 76 proteins were uniquely expressed in the pEVs-treated group, while 49 proteins were exclusively identified in the PBS control group. These unique proteins were classified according to their relevance to neurogenesis-promoting functions and presented as percentages based on the number of proteins in each category.

| **Category** | **pEVs: DG unique 76** | | **PBS: DG unique 49** | |
|---|---|---|---|---|
| | **Count** | **Percentage** | **Count** | **Percentage** |
| Signaling/Growth Factors | 21 | 27.60% | 16 | 32.60% |
| Transcription/Nuclear | 20 | 26.30% | 7 | 14.30% |
| Transport/Carrier | 10 | 13.20% | 15 | 30.60% |
| Enzymes/Metabolism | 9 | 11.80% | 2 | 4.10% |
| Structural/Cytoskeletal | 5 | 6.60% | 2 | 4.10% |
| Apoptosis/Cell Death | 3 | 4.00% | 0 | 0.00% |
| Cell Adhesion/ECM | 2 | 2.60% | 1 | 2.00% |
| Chaperones/Folding | 2 | 2.60% | 1 | 2.00% |
| Ion Transport | 0 | 0.00% | 3 | 6.10% |
| Other/Unknown | 3 | 4.00% | 0 | 0.00% |
| Lipid Metabolism | 0 | 0.00% | 1 | 2.00% |
| Proteasome/Degradation | 1 | 1.30% | 0 | 0.00% |
| Redox/Stress Response | 0 | 0.00% | 1 | 2.00% |

**Supplemental Table S3. Top 10 upregulated and downregulated differentially expressed proteins (DEPs) in the DG region following pEVs treatment.** The table presents the top 10 upregulated and downregulated differentially expressed proteins (DEPs) in the DG region after pEVs treatment compared to the PBS control group. Protein expression changes were identified through DEP analysis using a fold-change threshold of ≥ 1.2 and a significance cutoff of $p \leq 0.05$. These DEPs highlight the molecular alterations induced by pEVs treatment.

| | | | |
|---|---|---|---|
| **Top 10 upregulated** | | | |
| 1 | P02088 | Log2FC:3.38 | Hemoglobin subunit beta-1 |
| 2 | P01942 | Log2FC:2.71 | Hemoglobin subunit alpha |
| 3 | Q3UHD1 | Log2FC:2.13 | Adhesion G protein-coupled receptor B1 |
| 4 | P09470 | Log2FC:1.65 | Angiotensin-converting enzyme |
| 5 | P13808 | Log2FC:1.35 | Anion exchange protein 2 |
| 6 | P32233 | Log2FC:1.22 | Developmentally-regulated GTP-binding protein 1 |
| 7 | P07724 | Log2FC:1.10 | Albumin |
| 8 | Q9JKC8 | Log2FC:1.09 | AP-3 complex subunit mu-1 |
| 9 | P97823 | Log2FC:0.99 | Acyl-protein thioesterase 1 |
| 10 | Q8BIK4 | Log2FC:9.96 | Dedicator of cytokinesis protein 9 |
| **Top 10 downregulated** | | | |
| 1 | Q8C5H8 | Log2FC: -1.30 | NAD kinase 2, mitochondrial |
| 2 | P63166 | Log2FC: -1.14 | Small ubiquitin-related modifier 1 |
| 3 | Q61599 | Log2FC: -0.90 | Rho GDP-dissociation inhibitor 2 |
| 4 | P56382 | Log2FC: -0.89 | ATP synthase subunit epsilon, mitochondrial |
| 5 | Q35083 | Log2FC: -0.89 | 1-acyl-sn-glycerol-3-phosphate acyltransferase alpha |
| 6 | Q8K4Z5 | Log2FC: -0.84 | Splicing factor 3A subunit 1 |
| 7 | Q8BFQ8 | Log2FC: -0.83 | Glutamine amidotransferase-like class 1 domain-containing protein 1 |
| 8 | Q62189 | Log2FC: -0.82 | U1 small nuclear ribonucleoprotein A |
| 9 | P10711 | Log2FC: -0.81 | Transcription elongation factor A protein 1 |
| 10 | Q3U0V2 | Log2FC: -0.78 | Tumor necrosis factor receptor type 1-associated DEATH domain protein |

**S**

### Supplemental Figures (Figures S1-S3)

**Supplemental Figure s1. Ex vivo imaging of neurosphere uptake of Alexa fluor 568-conjugated pEVs and HPPL** Internalization of Alexa Fluor 488-labeled pEVs and HPPL by neurospheres after 24-hour incubation, compared to untreated controls. Scale bar: 200 µm. Stellaris microscope with Z-stack to visualize 3D internalization within the neurospheres with (20x objectives).

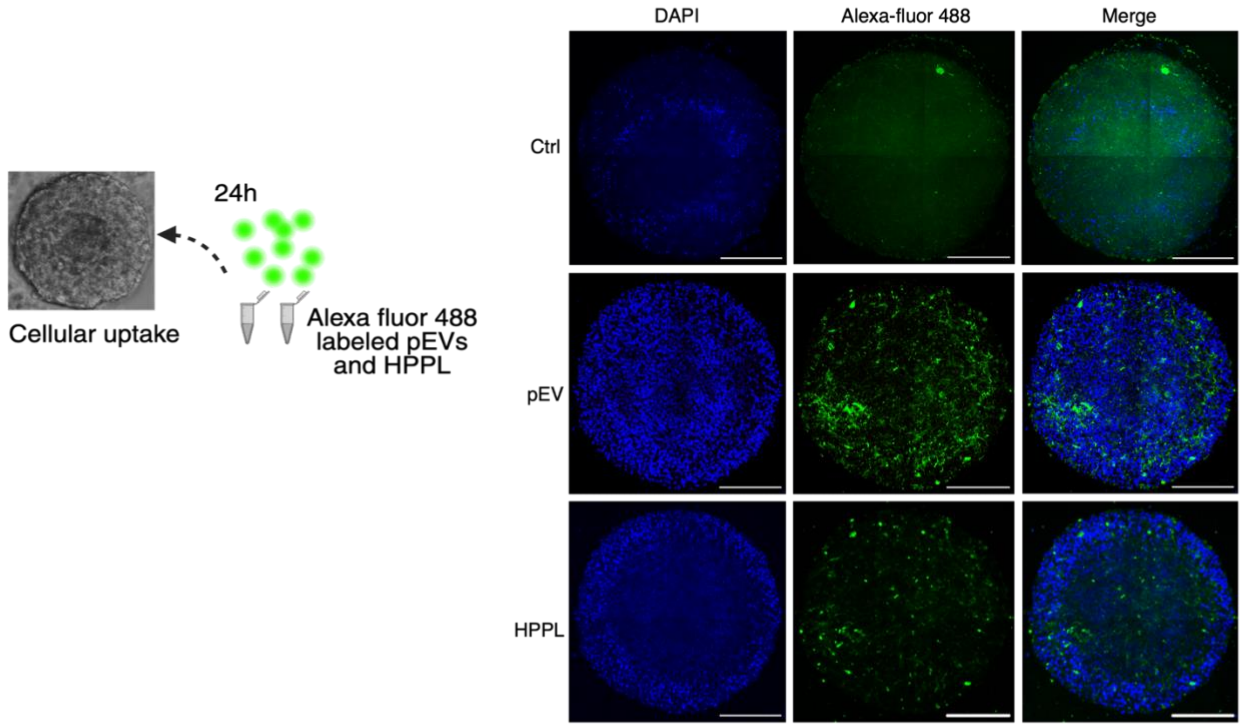

**Supplemental Figure s2. Gene Ontology (GO) analysis of uniquely expressed proteins in dentate gyrus tissue following pEVs or PBS treatment.** **A**. GO term enrichment analysis of proteins uniquely identified in the DG of pEVs-treated versus PBS-treated mice. Results are categorized by biological processes and molecular functions, highlighting treatment-specific pathways and molecular signatures associated with each condition.

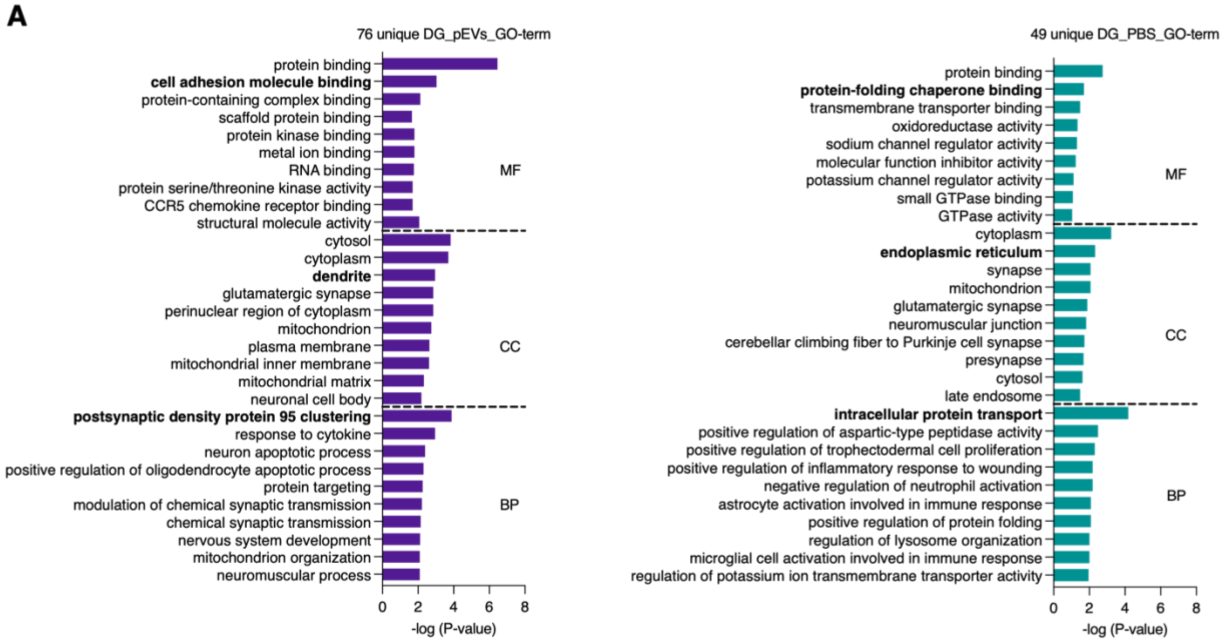
